# A small-molecule myosin inhibitor as a targeted multi-stage antimalarial

**DOI:** 10.1101/2022.09.09.507317

**Authors:** Darshan V. Trivedi, Anastasia Karabina, Gustave Bergnes, Alice Racca, Heba Wander, Seongwon Jung, Nimisha Mittal, Tonnie Huijs, Stephanie Ouchida, Paul V. Ruijgrok, Dan Song, Sergio Wittlin, Partha Mukherjee, Arnish Chakraborty, Elizabeth A. Winzeler, Jeremy N. Burrows, Benoît Laleu, Annamma Spudich, Kathleen Ruppel, Koen Dechering, Suman Nag, James A. Spudich

## Abstract

Malaria is a devastating disease that resulted in an estimated 627,000 deaths in 2020. About 80% of those deaths were among children under the age of five. Our approach is to develop small molecule inhibitors against cytoskeletal targets that are vital components of parasite function, essential at multiple stages of parasite infection, can be targeted with high specificity, and are highly druggable. Here we describe KNX-115, which inhibits purified *Plasmodium falciparum* myosin A (PfMyoA) actin-activated ATPase with a potency in the 10s of nanomolar range and >50-fold selectivity against cardiac, skeletal, and smooth muscle myosins. KNX-115 inhibits the blood and liver stages of *Plasmodium* with an EC_50_ of about 100 nanomolar, with negligible liver cell toxicity. In addition, KNX-115 inhibits sporozoite cell traversal and blocks the gametocyte to oocyst conversion in the mosquito. KNX-115 displays a similar killing profile to pyrimethamine and parasites are totally killed after 96 hours of treatment. In line with its novel mechanism of action, KNX-115 is equally effective at inhibiting a panel of *Plasmodium* strains resistant to experimental and marketed antimalarials. *In vitro* evolution data likely suggests a refractory potential of KNX-115 in developing parasite resistance.

## Introduction

Cells are composed of cytoskeletal networks of actin and microtubule tracks. These tracks are utilized by molecular motors like myosins, kinesins and dyneins to perform motile and force-generating processes in the cell [1]. This cytoskeletal system is vital to many essential cell functions in eukaryotes and is highly druggable, with multiple therapeutics currently in clinical trials [2-4] (NCT04823897, NCT04503265, NCT03856541), and one recently approved by the FDA for treatment of hypertrophic cardiomyopathy [5]. Kainomyx Inc. is a company devoted to using its extensive expertise in cytoskeletal biology for novel drug discoveries to treat diseases, with a primary focus on malaria.

In 2020, there were an estimated 241 million malaria cases worldwide, resulting in 627,000 deaths, and nearly 80% of those deaths occurred in children under the age of five [6]. Resistance to drugs within current treatment regimens, such as artemisinin and its derivatives, is increasing in Southeast Asia and Africa, and there is an urgent need to develop new therapeutics [7]. Many efforts for new drug discovery have focused on phenotypic screens to find small molecules that inhibit the invasion/multiplication of *Plasmodium* parasites in the blood stages of the parasite. The molecular targets of such screening hits may be unknown at the start of a project, although in many cases target-deconvolution methods, such as *in vitro* evolution and whole genome analysis, has identified the target [8]. Indeed, most of the newer and most advanced candidates in the global antimalarial drug development portfolio have come from this approach.

In general, however, knowing the target and understanding it well leads to a higher success rate in drug discovery.

Our approach is to develop small molecule inhibitors against the mechanistically well-understood cytoskeletal targets that are vital components of *Plasmodium* parasite function and are likely to be essential at multiple stages of parasite infection. According to gene deletion studies, three *Plasmodium falciparum* myosins, MyoA, MyoF and MyoK, have been shown to be independently essential for asexual blood stage parasite survival [9-11]. While all three of these myosins are potentially excellent targets for drug development, *Plasmodium falciparum* MyoA (PfMyoA), which belongs to the apicomplexan-specific class XIV family of myosins, was of primary interest for us to pursue as a first target because of its essential role in host cell invasion [12, 13]. This single-headed myosin motor powers the motility of the sporozoite, merozoite and ookinete stages of the parasite [12-15] through its ATP-dependent cyclical interactions with actin in the glideosome, a macromolecular complex necessary for the parasite’s motility and host cell invasion [16]. It is also biochemically well-characterized by *in vitro* studies using recombinantly expressed full-length Pf MyoA [17]. These studies revealed that motor function can be modulated by phosphorylation to optimize force and velocity parameters for the motile and invasion functions of the parasite during its human and mosquito life cycles [12]. The high-resolution structure of PfMyoA combined with biochemical and parasitological studies [14, 15] has led to a detailed understanding of how this molecular motor works [12, 18-20].

We have leveraged the amenability of PfMyoA to such powerful molecular, structural, and cellular drug discovery tools to identify KNX-115, a 17-nM inhibitor of the PfMyoA ATPase. KNX-115 results from structure-activity-relationship (SAR) studies on the original sub-micromolar-affinity hit KNX-002, obtained in a screen carried out by Cytokinetics Inc, and licensed by Kainomyx. We demonstrate that by inhibiting PfMyoA, KNX-115 inhibits multiple stages of the *Plasmodium* life cycle, an attractive feature for a malaria therapeutic. In addition, in the companion bioRxiv papers, the laboratories of Dr. Kathleen Trybus and Dr. Anne Houdusse have joined forces to describe biochemical measurements of the actomyosin cell cycle and the structure of PfMyoA with KNX-002 bound, and the Dr. Gary Ward laboratory shows that KNX-002 inhibits *Toxoplasma* invasion.

## Results

### KNX-115 inhibits PfMyoA using *in vitro* molecular assays and inhibits multiple steps in the parasite life cycle using *in vitro* cellular assays

#### KNX-115 inhibits actin-activated PfMyoA ATPase with nanomolar potency

KNX-002, the original hit licensed from Cytokinetics, showed inhibition of full-length PfMyoA in an actin-activated myosin ATPase assay with an IC_50_ of 0.4 ± 0.1 µM (n = 5; see Figure 1A for a representative experiment). SAR on KNX-002 led to KNX-115, with an IC_50_ of <40 nM (Figure 1A). Since this measured IC_50_ is very close to the PfMyoA concentration used in the assay (40 nM), we used the Morrison equation [21], which does not assume that the free concentration of the inhibitor equals the total concentration, to estimate the inhibition constant, K_i_, to be 17 nM (Figure 1B), a >20-fold gain in potency over KNX-002.

**Figure 1:**
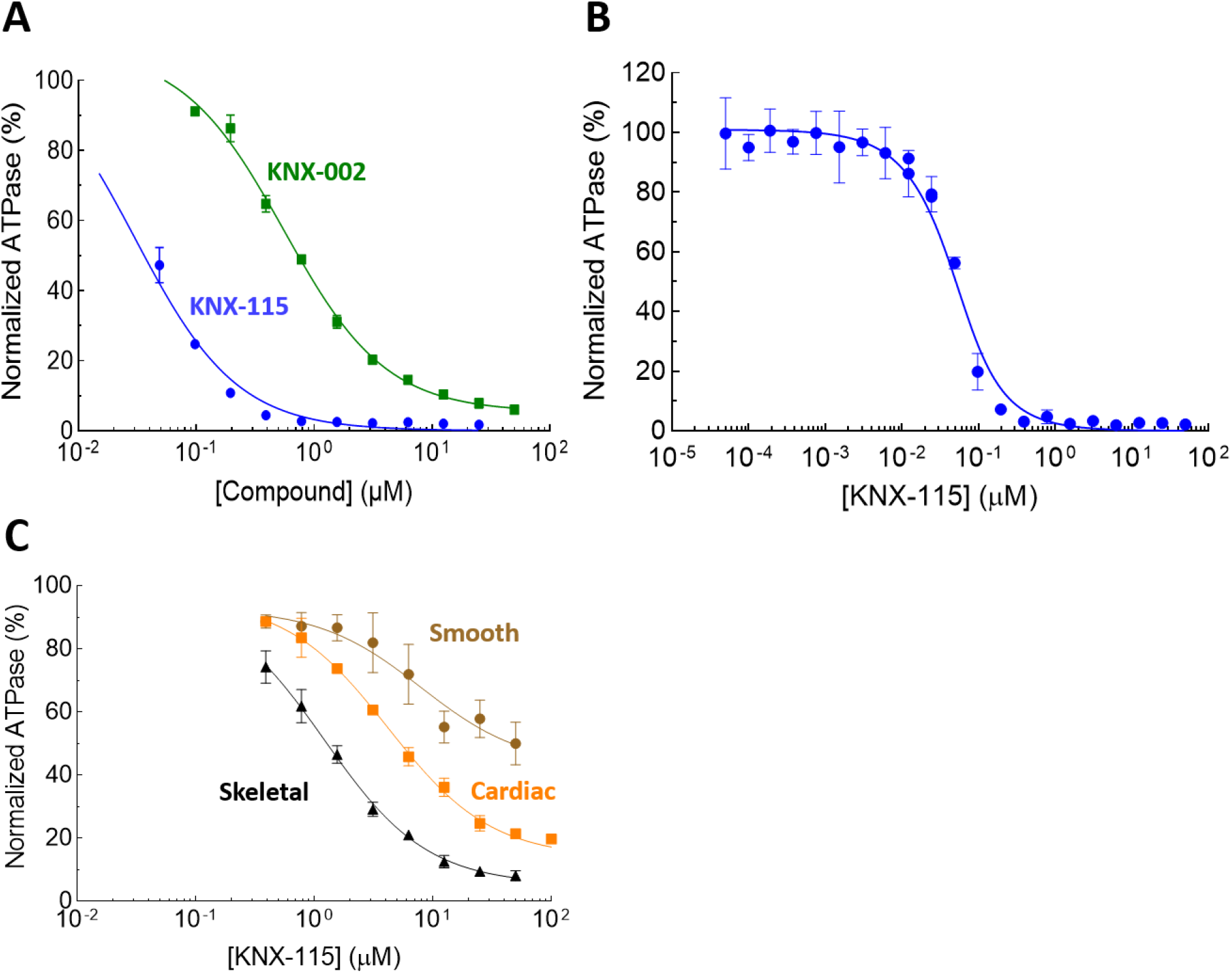
KNX-115 potently and selectively inhibits PfMyoA. (A) Actin-activated PfMyoA ATPase inhibition by parent compound KNX-002 (green) and its derivative KNX-115 (blue). KNX-002 IC_50_ = 0.6 µM and KNX-115 IC_50_ < 0.04 µM. (B) Morrison fit of a 22-point curve of PfMyoA inhibition. K_i_ = 17 nM. (C) KNX-115 mediated inhibition of rabbit skeletal myosin, bovine cardiac myosin, and chicken smooth muscle myosin actin-activated ATPase activity with IC_50_ of 1 ± 0.3 µM, 4.2 ± 0.05 µM, and >40 µM, respectively. IC_50_ values are over 50-fold greater for all three muscle myosins, suggesting high selectivity for PfMyoA. 100% activity is the actin-activated ATPase rate in the DMSO control (ranges from 10-12 s^-1^, at 15 µM actin). IC50 is the concentration of KNX-115 needed to reduce the normalized ATPase rate to 50%. Data are expressed as mean ± SD.

Dose-response curves of KNX-115 inhibition of bovine cardiac, rabbit skeletal, and chicken smooth muscle myosins showed IC_50s_ of 10 ± 6.4 µM (n = 3), 1 ± 0.4 µM (n = 6), and >50 µM (n = 2), respectively (see Figure 1C for representative experiments). Considering a K_i_ for PfMyoA of 17 nM, selectivity is greater than 50-fold for all three muscle myosins, suggesting high specificity for PfMyoA.

#### KNX-115 inhibits parasite growth in both blood and liver stages of Plasmodium

We next tested the hypothesis that inhibiting PfMyoA blocks the growth of *Plasmodium* parasites in the asexual blood stage. Inhibition could potentially involve any of multiple steps in the parasite asexual reproductive cycle, including invasion of the red blood cells, development and schizogony of the parasites in the red blood cell after invasion, and egression from the red blood cell [12, 13, 15, 22]. In asynchronous asexual blood stage cultures, KNX-115 inhibited the growth of *P. falciparum* in red blood cell cultures with an effective concentration (EC_50_) of 162 ± 14 nM (SEM, n = 52) (Figure 2A). This weakening of KNX-115 potency in parasite cellular assays versus the biochemical assays is in line with weakening of KNX-115’s biochemical potency in the presence of Albumax (Figure S2). Albumax, a media additive in parasite cellular assays, binds small molecules and leads to a decrease in the free fraction of the small molecule thus leading to apparent weakening of potencies in engaging the target.

**Figure 2:**
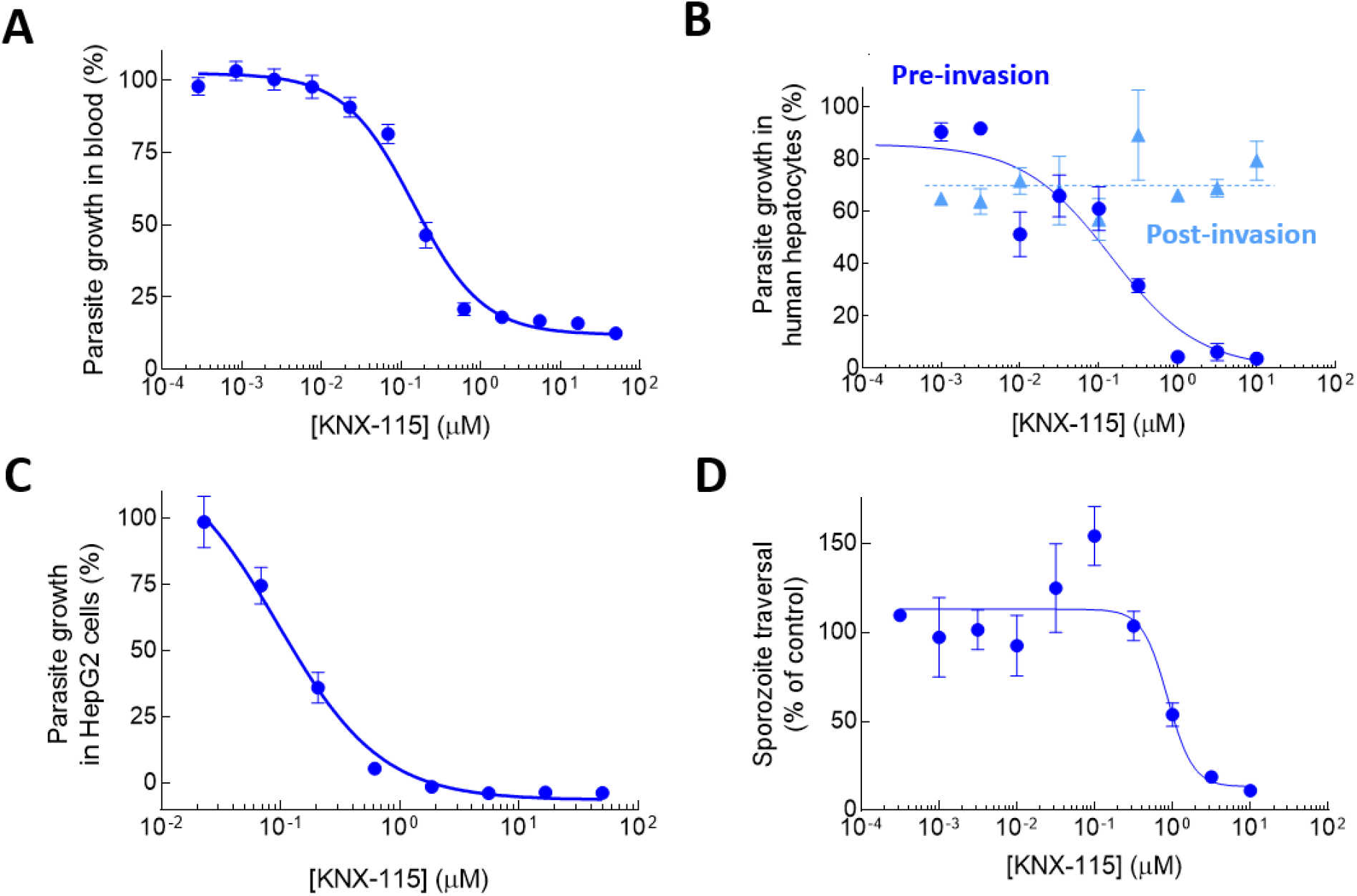
KNX-115 inhibits the asexual blood stage replication, liver stage invasion, and sporozoite cell traversal in *Plasmodium* parasites. (A) KNX-115 dose-dependently inhibits the growth of *P. falciparum* in an asexual blood stage assay with an EC50 of 162 ± 14 nM. (B) KNX-115 dose-dependently inhibits the growth of *P*. f*alciparum* in a liver stage assay using human hepatocytes when sporozoites are preincubated with KNX-115 (blue) with an EC_50_ of 182 ± 64 nM. The inhibition is lost when KNX-115 is added post-invasion (light blue). Data are from two biological replicates. Error bars indicate standard deviations. (C) KNX-115 dose-dependently inhibits the growth of *P. berghei* in a liver stage development assay using HepG2 cells with an EC_50_ of 108 ± 21 nM [25]. Data are from two experiments performed on two different days. (D) KNX-115 inhibits sporozoite cell traversal through HC-04 cells with an EC_50_ of 816 ± 55 nM. Data are from two biological and 6 technical replicates and are expressed as mean ± SEM.

A highly desirable feature of antimalarials is activity against multiple stages of the parasite life cycle. Given the importance of the glideosome in sporozoite motility [14], we tested whether PfMyoA inhibition would affect the ability of the parasites to invade and multiply in liver cells, the first intracellular niche occupied by the parasite in a human host. When *P. falciparum* parasites were preincubated with KNX-115 before adding them to primary human hepatocytes, inhibition of parasite multiplication in the hepatocyte culture was observed with an EC_50_ of 182 nM ± 64 nM (n = 2, see Figure 2B for a representative experiment), essentially the same level of inhibition as was seen in the asexual blood stage assay. Nearly identical results were obtained in experiments measuring inhibition of multiplication of *P. berghei* parasites in HepG2 cells when compounds were added preinvasion (EC_50_ = 108 ± 21 nM, SEM, n = 21) (Figure 2C). Interestingly, when *P. falciparum* parasite invasion was allowed to occur before adding KNX-115, no inhibition of parasite multiplication was seen (Figure 2B), suggesting that KNX-115 has its most potent activity early in infection, possibly by blocking invasion. We cannot rule out, however, potential metabolism of KNX-115 by human hepatocytes that might prevent inhibition of parasite function inside these cells. Importantly, no liver cell toxicity was seen in the human hepatocyte experiments even at 10 µM KNX-115 (Figure S1B), and no evidence of toxicity was observed below KNX-115 concentrations of 5 µM in the HepG2 cells (Figure S1A).

#### KNX-115 inhibits sporozoite cell traversal

An essential part of the parasite life cycle is the first step of cell traversal of sporozoites from the site of their injection in the dermis by a mosquito to the liver by gliding motility and transmigration through cells. It is rare to find an inhibitor of sporozoite cell traversal, but we hypothesized that we would observe such inhibition by explicitly targeting PfMyoA, which is a pivotal component of the glideosome used for sporozoite motility. Indeed, KNX-115 inhibited *P. falciparum* sporozoite cell traversal with an EC_50_ of 816 ± 55 nM (n = 2) (Figure 2D).

#### KNX-115 inhibits the gametocyte to oocyst conversion in the mosquito

Our findings that inhibiting PfMyoA blocks parasite development in both blood cells and liver cells and inhibits sporozoite cell traversal indicate the importance of PfMyoA in many fundamental aspects of the parasite life cycle. The mosquito stages of malaria parasites start with the activation of gametogenesis from the intraerythrocytic gametocytes immediately after a blood meal and end with the release of sporozoites. Given that the development of sporozoites from gametocytes in the mosquito involves a motile ookinete stage, we suspected a role of the cytoskeleton proteins and postulated that inhibiting PfMyoA would affect one or more of these stages. Consistent with our hypothesis, KNX-115 blocked the gametocyte to oocyst conversion in the mosquito host with an EC_50_ of 230 nM (Figure 3A). Unlike the dihydroartemisinin (DHA) control, KNX-115, did not block exflagellation events of male gametocytes (Figure 3B), in line with a lack of MyoA protein expression in the activated gametocyte stages of the parasite [10]. Similarly, MyoA protein expression is not detected in the non-activated gametocyte stages [10] and KNX-115 did not block the growth of stage III-IV gametocytes measured by a gametocyte viability assay (Figure 3C). The combined data indicate that the block in transmission is not through a gametocytocidal effect but occurs downstream of the intraerythrocytic gametocyte, presumably driven by inhibition of the motile ookinete stage of the parasite.

**Figure 3:**
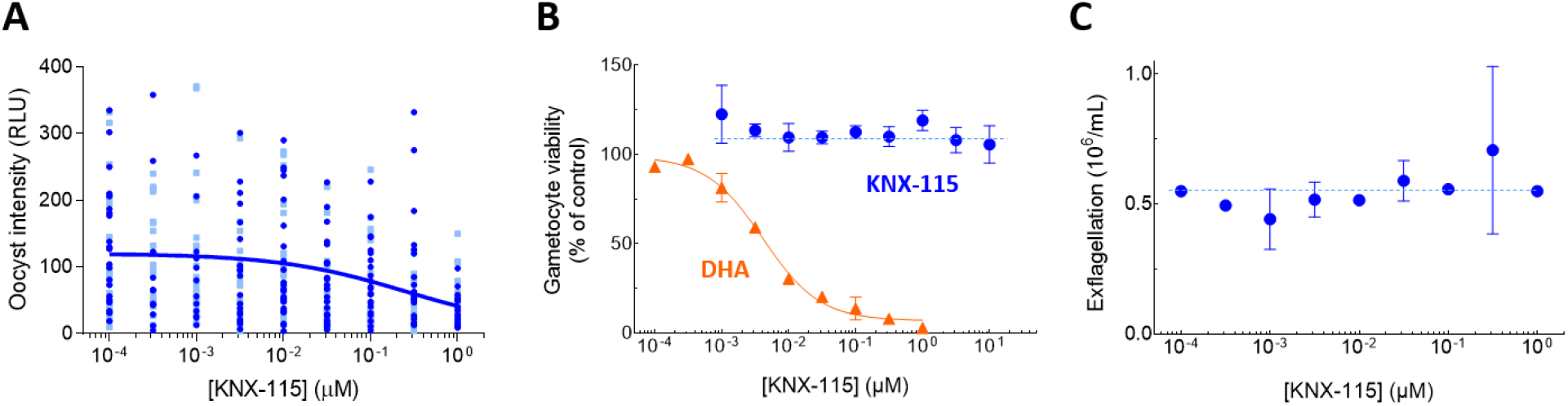
KNX-115 inhibits gametocyte to oocyst conversion in the mosquito and has no effect on the gametocyte viability and exflagellation capacity of the male gametocyte. A) KNX-115 dose-dependently inhibits the conversion of gametocytes to oocysts with an EC_50_ of 230 nM in a standard membrane feeding assay performed with a *P. falciparum* NF54-HGL reporter strain. Each point is oocyst intensity in individual mosquitoes from two independent feeds per drug concentration tested. B) KNX-115 is not gametocytocidal in a gametocyte viability assay. Dihydroartemisinin (DHA) inhibition is a positive control. C) KNX-115 does not affect exflagellation events in male gametocytes. Data are expressed as mean ± SD.

### KNX-115 is parasiticidal, acts on resistant strains, and selects resistance, but with mild reduction in potency

#### Parasite reduction ratio assays show that KNX-115 kills parasites and a stage-specificity assay shows that KNX-115 treated parasites cannot initiate a new round of parasite multiplication

While the in-vitro cellular assays show that KNX-115 inhibits the accumulation of parasites in red blood cell cultures and liver cells, it is important to determine whether such inhibitory effects are due to blocking parasite growth or killing them. This is determined by parasite reduction ratio (PRR) assays. The killing profile of KNX-115 at 5.4 µM showed that parasites were eliminated after 96 hours (Figure 4A).

**Figure 4:**
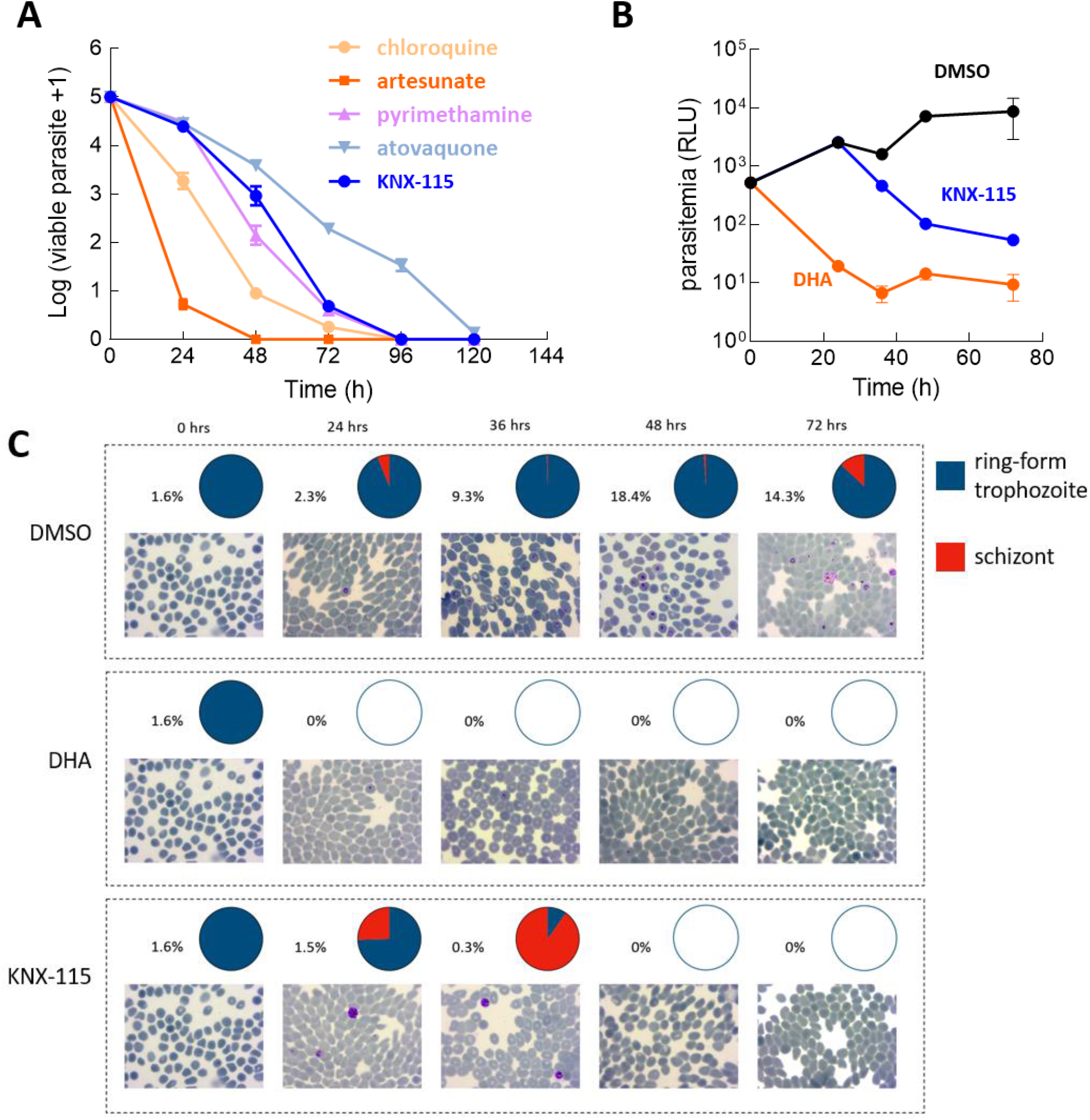
KNX-115 is parasiticidal and has a killing rate profile comparable to some existing antimalarials. (A) *P. falciparum* viability time-course profiles demonstrate KNX-115 (blue) is parasiticidal and viable parasites are killed by 96 hours. There is a lag phase of 24 hours. KNX-115’s killing kinetics are similar to pyrimethamine. The assay was performed at 10X IC_50_ of each compound. Data are expressed as mean ± SD. (B) Parasites of a nanoluc reporter strain were synchronized at the ring stage, exposed to 5 µM KNX-115, and luciferase activity was compared over time to that of control (DMSO or 1 µM DHA-treated) cultures. (C) Giemsa staining and parasite counting of cultures treated with KNX-115, vehicle control (0.1% DMSO) or 1 µM DHA. The percentage values are total parasitemia and the pie charts indicate the proportion of ring/trophozoite versus schizont stages. Data for vehicle controls and DHA-treated cultures are from three independent flasks, and data for KNX-115 are from a single flask.

Compared with other antimalarials, the killing profile closely resembled pyrimethamine, though slightly slower between 24 and 72 hours. Both showed a 24-hour lag before killing. This was faster than atovaquone but slower than chloroquine. Artesunate was faster than chloroquine, as is characteristic of this family of inhibitors (Figure 4A). Table 1 outlines the killing parameters of KNX-115; 99.9% of the parasite’s initial population was cleared in 57.8 hours.

**Table 1:** Killing curve parameters of KNX-115 compared to standard antimalarials pyrimethamine and chloroquine.

| Compound | Dose (μM) | Lag phase (h) | Slope | R<sup>2</sup> | Log PRR | PCT<sub>99.9%</sub> (h) |
| --- | --- | --- | --- | --- | --- | --- |
| KNX-115 | 5.35 | 24 | -0.0772 ± 0.005 | 0.959 | 3.70 | 57.8 |
| Controls |  |  |  |  |  |  |
| Pyrimethamine | 0.27 | 24 | -0.0811 ± 0.005 | 0.969 | 3.89 | 50.5 |
| Chloroquine | 0.22 | 0 | -0.0843 ± 0.004 | 0.980 | 4.05 | 36.8 |

In stage-specificity assays, cultures from a *P. falciparum* NF54 strain expressing a Nanoluciferase reporter were synchronized at the ring/trophozoite stage, treated with 5 µM KNX-115 and followed in time. The luminescence signal increased in the first 24 hours, comparable to levels in vehicle-treated cultures, and then dropped significantly (Figure 4B). In contrast, luminescence in DHA-treated cultures dropped immediately and did not show a 24-hour lag phase. Microscopy of the same cultures indicated a sharp increase in parasitemia in vehicle-treated cultures from 24 hours onwards (Figure 4C). DHA led to rapid killing of the parasites. KNX-115 treated parasites did not replicate in the first 24 hours and then showed a drop in parasitemia. The remaining parasites after 24 hours were mostly in the schizont stage. The combined data demonstrate that KNX-115 treated parasites are unable to initiate a new round of parasite multiplication.

#### KNX-115 is effective at inhibiting Plasmodium strains that are resistant to approved antimalarials and advanced portfolio molecules

Given the emergence of resistance to current antimalarials, a highly desirable profile of a new antimalarial is that it remains inhibitory against strains already resistant to approved antimalarials and advanced portfolio molecules. KNX-115 dose-response curves were essentially the same using Pf NF54 parasite strains showing multi drug resistance to chloroquine cycloguanil/pyrimethamine (strains 7GA and K1), chloroquine cycloguanil/pyrimethamine6, artemisinin and piperaquine (strain PH1263-C), and cycloguanil/pyrimethamine and atovaquone (strain TM90C2B), when compared to the parent Pf NF54 strain (Figure 5A). We also tested five Pf Dd2 strains each resistant to a different advanced portfolio compound in addition to chloroquine cycloguanil/pyrimetamine. Again, KNX-115 dose-response curves were essentially the same using the five resistant Pf Dd2 strains compared to the parent Pf Dd2 strain (Figure 5B), suggesting that KNX-115 is not subject to resistant mechanisms mediated through the genetic alterations represented by these parasite lines.

**Figure 5:**
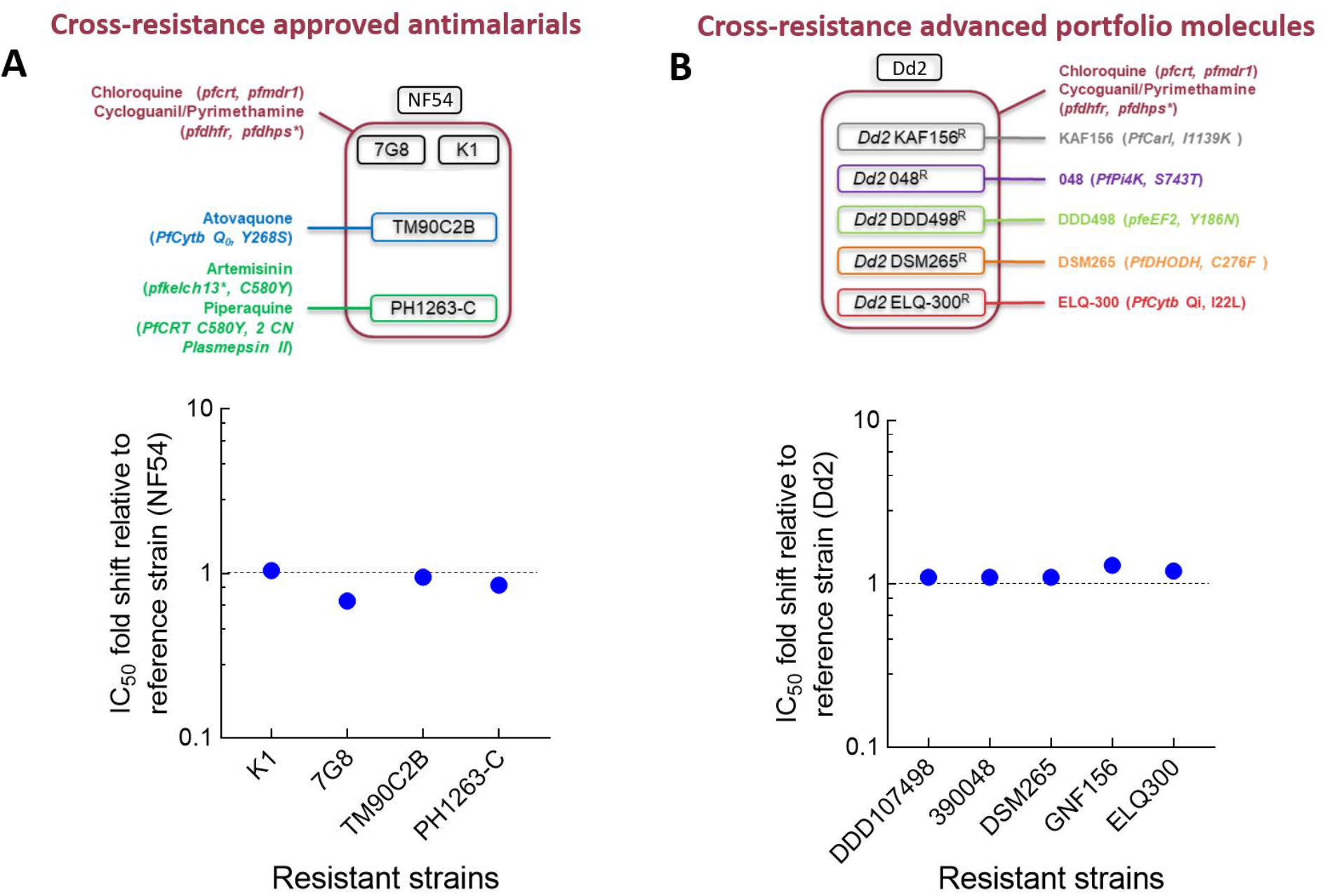
KNX-115 inhibits asexual blood stage parasite replication in resistant strains of *P. falciparum*. (A) Top panel: Strains that are resistant to approved antimalarials. Five genetic backgrounds are represented in this sub-panel (7G8 – South America; TM90C2B, K1 – South-East Asia; NF54 West Africa). NF54 is considered the reference pan-sensitive strain and TM90C2B, PH1263-C, 7G8, K1 are multi-drug resistant (MDR) strains, in which seven drug resistant genes are represented. The star (*) indicates a resistance mechanism that cannot be detected performing a standard ABS proliferation assay Bottom panel: KNX-115 is equipotent in inhibiting asexual blood stage parasitemia in resistant strains of *P. falciparum* compared to the parental NF54 strain. (B) Top panel: Dd2 strains that are resistant to advanced portfolio molecules. Five genetically engineered strains in a Dd2 genetic background are present in this sub-panel, in which five drug resistance-correlated genes are represented. The parental Dd2 wild-type strain is used to determine a potential EC50 shift. Bottom panel: KNX-115 is equipotent in inhibiting asexual blood stage parasitemia in resistant strains of *P. falciparum* compared to the parental Dd2 strain. Data are expressed as mean from at least 2 independent biological replicates.

#### In vitro evolution of drug resistance shows resistant parasites that are still sensitive to KNX-115

Although KNX-115 proved potent against already resistant strains, we next checked the propensity of KNX-115 to induce resistance in parasites. Treatment of 10^8^ *P. falciparum* parasites (NF54 asynchronous cultures) with 3X EC_50_ of KNX-115 showed significant inhibition of parasite multiplication by four days, but recrudescent parasites were observed between 8-14 days into the experiment (Figure 6A). When three of these resistant clones were analyzed by dose-response curves using KNX-115, their EC_50s_ of inhibition were only marginally shifted to higher concentrations (Figure 6B). Thus, while the WT clone showed inhibition at an EC_50_ of 0.6 µM KNX-115, the three resistant strains showed inhibition at 2.2 µM, 2 µM and 1.2 µM KNX-115. This demonstrates a mild weakening of potency by ∼3 fold.

**Figure 6:**
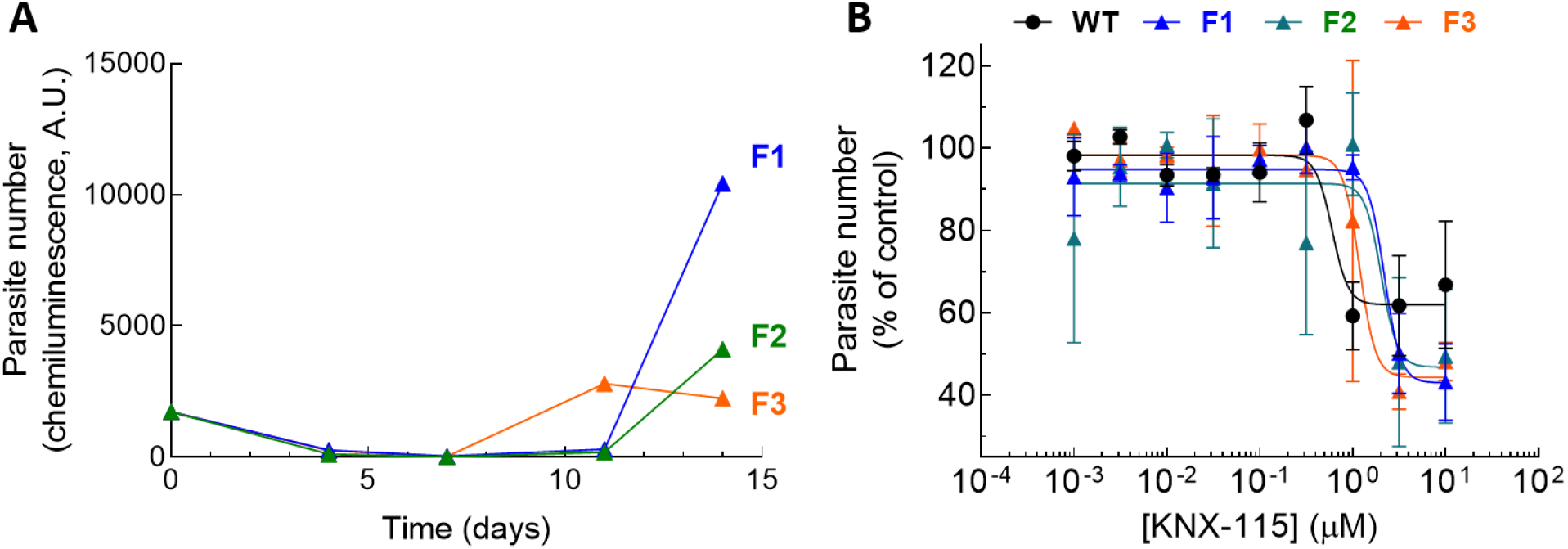
In vitro evolution of drug resistance shows mildly resistant parasites that are still sensitive to KNX-115. (A) Three flasks showing emergence of recrudescent *P. falciparum* parasites upon treatment with KNX-115 at 3X IC_50_. (B) KNX-115 dose-response curves with the recrudescent resistant parasites compared to the WT strain. The WT strain showed inhibition at an EC_50_ of 0.6 µM KNX-115, and the three resistant strains showed inhibition at 2.2 µM, 2 µM and 1.2 µM KNX-115. Data are from two replicates.

### Discussion and perspectives

Malaria is one of the most severe global health burdens leading to significant morbidity and mortality, primarily in young children and pregnant women in developing countries. Multiple studies have confirmed that malaria parasites in Africa and Southeast Asia have developed or are at risk of developing resistance to artemisinins - a family of widely used and critically important antimalarials. The increasing spread of artemisinin-resistant parasites and the failure of drugs such as piperaquine in SE Asia have served as major wake-up calls for the need to develop the next generation of robust therapies.

Using a well-tested approach to target mechanistically well-understood cytoskeletal targets, at Kainomyx we developed KNX-115 as a nanomolar potent small-molecule inhibitor of PfMyoA, an essential protein needed for motility and force-production in the various stages of the parasite life cycle (Figure 7). Here we have shown that KNX-115 inhibits parasite growth in the blood and liver cells, possibly due to a blockage in the parasite’s invasion ability. Moreover, the finding that accumulation of *P. berghei* and *P. falciparum* in liver cell cultures was inhibited to similar extents by KNX-115 suggests that inhibition of PfMyoA will likely be effective across *Plasmodium* species. This is a highly desirable property of a malaria therapeutic.

**Figure 7:**
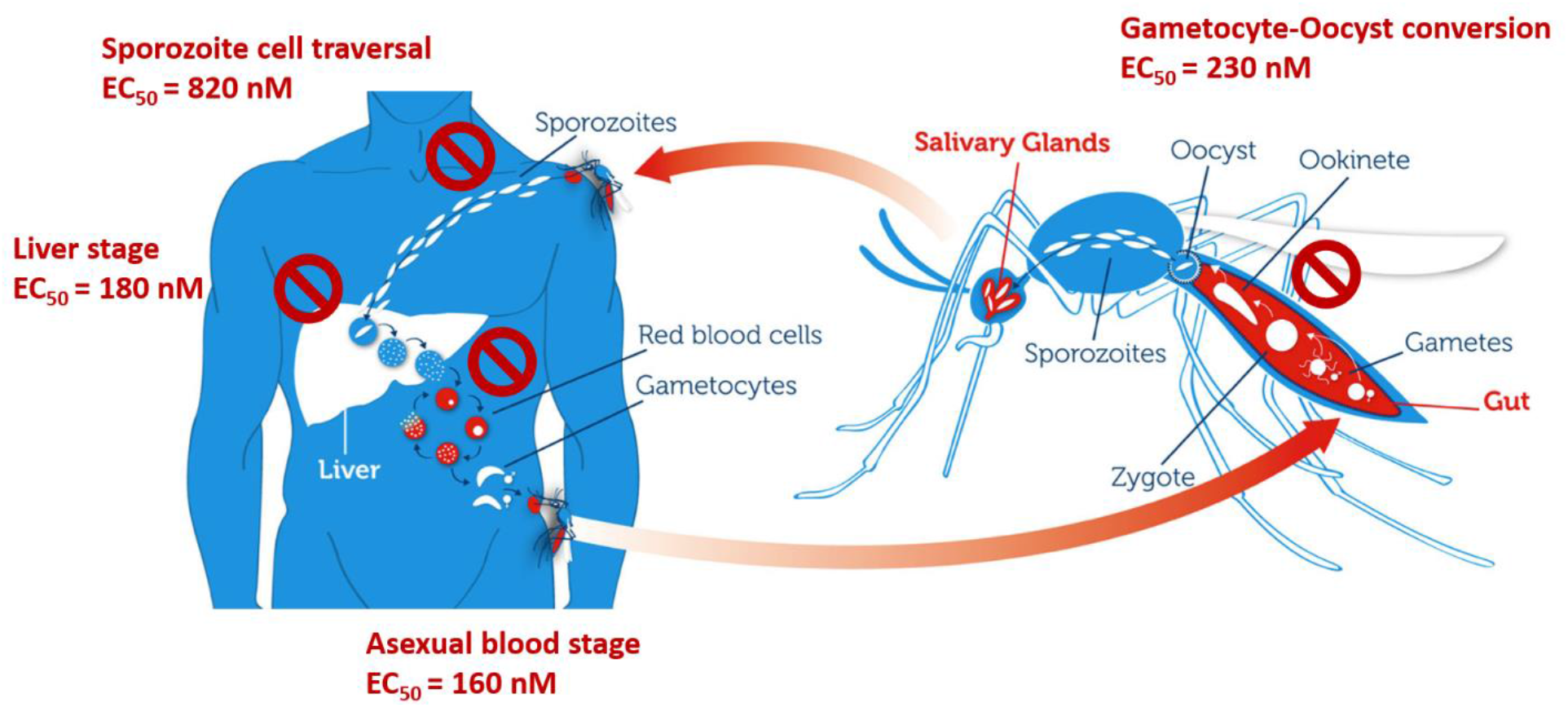
KNX-115 is a multistage inhibitor of the *Plasmodium* lifecycle. KNX-115 inhibits the asexual blood stage, liver stage and sporozoite cell traversal in human host. It also inhibits the parasite development in the mosquito midgut. This figure was modified from the original published in [26].

Since no inhibition of parasite growth was observed when parasites were allowed to invade cultured human hepatocytes prior to treatment with KNX-115, this molecule appears to inhibit the invasion process rather than affecting the development of schizonts within the liver cells. One cannot rule out, however, that KNX-115 also inhibits the development and growth of parasites in the liver cells. The lack of inhibition by KNX-115 post-invasion could reflect, for example, a permeability problem or metabolic clearance of KNX-115 before it had a chance to act on its target. More experiments are necessary to answer this question.

In line with the observed inhibition of hepatocyte invasion, KNX-115 inhibits sporozoite cell traversal. This suggests that KNX-115 could act as a prophylactic agent, provided the inhibition kinetics are fast enough to prevent human hepatocyte infection. Similarly, KNX-115 blocks the gametocyte to oocyst conversion which involves a motile zygote stage. Both sporozoites and ookinetes use a MyoA-containing glideosome protein structure to exert gliding motility [14]. More recently it has been shown that blood stage merozoites display gliding motility [15]. In light of our observation that KNX-115 prevented initiation of a new cycle of asexual blood stage replication, it is tempting to speculate that KNX-115 inhibits merozoite motility and thereby prevents red blood cell invasion. In line with these observations, KNX-115 did not inhibit the exflagellation of male gametocytes or the development of stage III-IV gametocytes which likely do not involve any gliding functions required by the parasite. This is consistent with a report that shows that MyoA protein is not expressed in the non-activated and activated gametocyte stages of *P. berghei* [10].

KNX-115 inhibits the development of the parasite in the mosquito midgut by likely inhibiting the motile ookinete stage of the parasite. This property of MyoA inhibitors can be leveraged to block parasite transmission in a dual manner. Anopheles populations are growing resistant to insecticides (applied to bed nets) leading to a global threat of resurgence of malaria. Paton et al [23] have demonstrated that mosquitoes can take up the transmission-blocking antimalarial atovaquone from treated surfaces. This causes full parasite arrest in the mosquito midgut and prevents transmission. Provided a similar favorable contact-dependent uptake, MyoA inhibitors like KNX-115 can likely be utilized for impregnating mosquito nets that can significantly mitigate insecticide resistance in these mosquitoes while blocking transmission of the parasite. Modeling studies have identified such efforts as promising leads for eradicating malaria [23, 24]. In addition, a MyoA inhibitor designed with an ideal pharmacokinetic profile could be taken up by an Anopheles mosquito in a blood meal which could in turn lead to the blocking of parasite development in the mosquito midgut. This could also lead to malaria control by blocking transmission.

KNX-115 shows a lag of 24 hours before it demonstrates killing in the PRR and stage-specificity assays. This might be due to an asynchronous starting culture with >80% rings in the PRR assay. If KNX-115 prevents RBC re-invasion, then in a truly synchronized culture, addition of KNX-115 to the culture primarily in early ring stage would result in a 48-hr lag. Similarly, if the compound were added at the end of schizogony, a 0-h lag would be observed. In a clinical setting where patients would likely have a mixture of different stages of the parasite, a 24-h lag before killing is a reasonable assumption. Given its slower killing rate, a MyoA candidate drug will likely need to be combined with a faster acting drug. In the stage-specificity assay, KNX-115 leads to stagnation of parasitemia in the first 24 hours and is accompanied by a sharp increase in the schizont population relative to ring stage population. This is followed by killing of the parasites. This indicates that KNX-115 likely works by inhibiting invasion of merozoites released from the schizonts and possibly not inhibiting the ring/trophozoite stages. However, one cannot rule out the dual mechanism of egress-inhibition and invasion-inhibition by KNX-115. Earlier GAP45 knockout studies ruled out MyoA’s role in egress of merozoites from RBCs, but there can be significant phenotypic differences between a MyoA knockout parasite line and parasites with functionally inhibited MyoA.

The data that KNX-115 is equipotent in strains of parasites resistant to other antimalarials and advanced portfolio compounds is very encouraging and indicates that efficacy of a MyoA inhibitor is unlikely to be hampered by pre-existing resistance mechanisms. Our experiments concerning the generation of resistant species upon KNX-115 treatment suggest that the selected parasites display only a weak resistance to KNX-115, but more experiments are needed to uncover the resistance mechanisms of this inhibitor class on this cytoskeletal target. There can be an evolutionary fitness cost for the parasite to develop resistance by mutating MyoA to escape MyoA inhibitors.

In summary, the growing resistance of *Plasmodium falciparum* to existing antimalarials is a significant threat. This puts even more pressure on researchers and drug makers to find new viable treatments and/or vaccine options for malaria soon. Kainomyx is focused on contributing to this goal with its targeted therapeutic approach, which lies at the heart of modern drug discovery. Our discovery of KNX-115 puts forward a hypothesis of developing MyoA inhibitors as next generation of molecules for treating malaria.

## Acknowledgements

We thank Anna Adam as MMV project coordinator, Christian Scheurer (STPH) for his support with the *in vitro* assays of cross-resistance, and Rajib Barik and Subrata Chattopadhyay from TCGLS *in vitro* parasitology team for their support with the parasite reduction ratio assays. We would also like to thank Gang Liu from Bill & Melinda Gates Foundation for his comments on the manuscript. We thank Bill and Melinda Gates Foundation for support to carry this project forward. We are grateful to Cytokinetics for access to parent compounds that led to the development of KNX-115.

## Methods

### KNX-115

KNX-115 was derived from KNX-002 (formerly CK2140597), which was originally identified from high throughput screening of a library of 50,000 compounds performed at Cytokinetics, to identify inhibitors of the actin-activated ATPase of *T. gondii* MyoA (protein provided by Gary Ward, University of Vermont), and PfMyoA (protein provided by Kathleen Trybus, University of Vermont). Kainomyx licensed KNX-002 and 5 other compounds from Cytokinetics for drug discovery for malaria and other parasitic diseases in 2020.

### Protein expression and purification

PfMyoA was expressed as described previously [17] in Sf9 cells adapted to serum-free ESF 921 media purchased from Expression Systems. Sf9 cells were co-infected with PfMyoA, PUNC chaperone, PfELC and PfMTIP light chains, and harvested 48 hours post-infection and stored at -80°C. and purified as described before [17]. Briefly, PfMyoA was purified from Sf9 cell pellets that were resuspended in 25 mL lysis buffer (10 mM imidazole, pH 7.4, 0.2 M NaCl, 1 mM EGTA, 2.5 mM MgCl_2_, 7% (w/v) sucrose, 2 mM DTT, 0.5% (v/v) Igepal, 0.5 mM phenylmethanesulfonyl fluoride, 5 μg/ml leupeptin, 5 μg/ml pepstatin A, 1X Roche cOmplete EDTA free tablet, 2 mM MgATP) per billion cells and lysed by dounce homogenization. An extra 2.5 mM MgATP was added before clarification at 266,000 x g for 40 min. The supernatant was passed through a 1.2 µM filter before batch binding to ANTI-FLAG M2 affinity resin (Sigma) for 1 hour. Resin was packed into a chromatography column and washed with 10 volumes of 10 mM imidazole, pH 7.4, 0.2 M NaCl, 1 mM EGTA, 0.5 mM phenylmethanesulfonyl fluoride. This was followed by 5 wash volumes of 10 mM imidazole, pH 7.4, 5 mM KCl, 3 mM MgCl_2_, 1 mM DTT, and PfMyoA was then eluted in the same buffer supplemented with 0.1 mg/mL FLAG peptide. Fractions containing myosin were pooled, washed, and concentrated using a 10,000 MWCO Amicon centrifugal filter device (Millipore) in 10 mM imidazole, pH 7.4, 5 mM KCl, 3 mM MgCl_2_, 10% (w/v) sucrose, 1 mM DTT, and stored at -80°C.

Skeletal and cardiac muscle myosins were purified as described previously [27], and smooth muscle myosin was purified as described before [28]. Bovine cardiac actin was purified as described previously [29].

### Myosin inhibition assay

The enzymatic activity of recombinantly expressed *Plasmodium falciparum* myosin A (MyoA) was assessed by the ATPase assay in the presence of several small molecule agents. This assay couples the release of adenosine diphosphate (ADP) from MyoA to the enzymes pyruvate kinase (PK) and lactate dehydrogenase (LDH) while monitoring the decrease in absorbance of nicotinamide adenine dinucleotide (NADH) at 340 nm as a function of time. MyoA hydrolyzes adenosine triphosphate (ATP) to ADP and phosphate. PK converts ADP back to ATP by converting phosphoenolpyruvate to pyruvate. This pyruvate is then converted to lactate by LDH by converting NADH (nicotinamide adenine dinucleotide) to NAD (oxidized nicotinamide adenine dinucleotide). MyoA was recombinantly expressed in the baculovirus/Sf9 insect cell expression system. Assay was performed in 50 µL reaction volume in a 96 well half area plate. Final assay conditions were 50 nM of MyoA, 10 mM Imidazole, pH 7.4, 5 mM KCl, 3 mM MgCl_2_, 1 mM DTT, 2 mM ATP, 1.5 mM PEP, 1 mM NADH, 20 units/mL LDH,100 units/mL PK.

Stock solutions of the compounds were made in 100% DMSO and a dilution series of the compound was created in DMSO. This dilution series was created such that the final required compound concentration in the assay would have a fixed DMSO concentration of 2% (v/v) in a 50 µL reaction volume. Duplicates of 8 or 12-point dose response curves were typically measured for each experiment. MyoA was incubated with the compound for two minutes and the reaction was started by addition of actin and coupling solution that contained ATP, PEP, NADH, LDH and PK. The decrease in NADH absorbance was followed for 30 minutes at 340 nm on a Molecular Devices M5e or Spectramax Plus 384 plate reader using clear half area plates. The ATPase rate at each compound concentration was normalized to the average ATPase rate at 2% DMSO. This normalized rate was plotted as a function of compound concentration and IC50 values (compound concentration at which the activity is inhibited by fifty percent) were determined by the CDD (Collaborative Drug Discovery) software. Any compound that did not achieve a fifty percent inhibition at the highest concentration tested was reported to have an IC50 greater than the highest concentration tested. Typically, all measurements were done at a highest compound concentration of 50 µM or 200 µM.

Selectivity measurements against skeletal, cardiac and smooth muscle myosin isoforms were performed as described above. Skeletal myosin was purified from rabbit muscle, cardiac myosin was purified from bovine ventricle and smooth muscle myosin was purified from chicken gizzard. The final concentrations of skeletal, cardiac and smooth muscle myosins in the ATPase assay were 100 nM, 200 nM and 300 nM respectively. Dose responses against these myosins were performed as described above.

### *Plasmodium falciparum* asexual blood stage assay

A dose response assay was performed to identify compounds that inhibit the growth of *P. falciparum* Dd2 strain parasites in human erythrocytes. 40 nL of compounds in 1:3 serial dilutions at 0.5% final DMSO concentration were transferred with an acoustic transfer system into black, clear-bottom plates. Artemisinin at a single concentration of 5 µM was used as a positive control, and 0.5% DMSO was used as a negative control. Parasite suspension with 0.3% parasitemia and 2.5% hematocrit in screening medium (RPMI 1640 with l-glutamine, without phenol red (Life Technologies, CA) supplemented with 0.2% AlbuMAX II lipid-rich BSA, 0.014 mg/mL hypoxanthine, 3.4 mM NaOH, 38.4 mM Hepes, 0.2% glucose, 0.2% sodium bicarbonate, and 0.05 mg/mL gentamicin) was prepared and dispensed into plates which already contained compounds to test. These plates were incubated for 72 h at 37C with water-soaked tissue in a bag gassed with 1% oxygen, 3% carbon dioxide and 96% nitrogen. After 72 h, SYBR Green I (Invitrogen) in Lysis buffer (20 mM Tris/HCl, 5 mM EDTA, 0.16% Saponin wt/vol, 1.6% Triton X vol/vol) was added and plates were incubated in the dark at room temperature for 24 h. After 24 h, fluorescence at 485 nm excitation and 530 nm emission was read from the bottom of the plates using the Perkin Elmer EnVision Multilabel Reader. The data was normalized as a percentage of the negative control (0.5% DMSO) and the background for the blood stage inhibition was defined as average fluorescence of the wells with Artemisinin at a single concentration of 5 µM. EC50 values were determined with Levenberg-Marquardt algorithm for curve fitting of the dose response data using CDD Vault software.

### *Plasmodium berghei* liver stage assay

For this assay, a human hepatoma cell line (HepG2-A16-CD81EGFP) was cultured at 37C in 5% C02 in DMEM media. 20-26 hours prior to sporozoite infection, the HepG2-A16-CD81EGFP cells were seeded in a white solid bottom plate. Thereafter, 18 hours prior to infection, 50 nL of compounds in 1:3 serial dilutions in DMSO (final 0.5% DMSO v/v) were transferred to the assay plates. Atovaquaone (5 µM) was used as a positive control and 0.5% DMSO was used as a negative control. *P. berghei* sporozoites were freshly obtained by dissecting salivary glands of infected *A. stephensi* mosquitoes and added to each well at a density of 1×103 sporozoites per well. The plates were centrifuged for 5 min at 330g and incubated at 37C for 48h in 5% C02 with high humidity. After 48h, the parasite growth was assessed by bioluminescence measurement. Media was removed by spinning the inverted plates at 150g for 30 seconds followed by addition of BrightGlo (Promega) for quantification of *P. berghei* parasites. Luminescence was measured by PerkinElmer Envision Multilabel Reader. EC50 values were determined in CDD vault normalized to maximum and minimum inhibition levels for the positive (Atovaquone 5 µM) and negative control (0.5% DMSO) wells.

### *Plasmodium falciparum* sporozoite cell traversal assay

Traversal assays were performed essentially as described before [30]. Human hepatoma HC-04 cells were seeded in 384-well plates in DMEM/F12 culture medium (Gibco) supplemented with 10% heat inactivated fetal calf serum (Gibco) and cultured at 37°C and 5% CO_2_ until 85-90% confluency. Test compounds were diluted in DMSO and then in culture medium to achieve a final DMSO concentration of 0.1%. Freshly isolated *P. falciparum* NF54 salivary gland sporozoites (SPZs) were pre-incubated with the compounds for 30 min at room temperature. Subsequently, fluorescent dextran was added to the serum-SPZ mix and the mix was transferred to the HC-04 cells. The plates were centrifuged and incubated for one hour at 37°C and 5% CO_2_. After incubation, the cells were washed with PBS-Tween and fixed with 4% paraformaldehyde. Hereafter, cells were counterstained with DAPI (4’,6-Diamidino-2-Phenylindole, Dihydrochloride) at room temperature. Cells were washed and the total number of cells and the number of traversed cells were quantified using an ImageXpress Pico automated cell imaging system (Molecular Devices).

### *Plasmodium falciparum* inhibition of liver stage development assay

Assessment of compound effects on *P. falciparum* NF54 invasion of human primary hepatocytes and intra-hepatocytic development was performed as described previously [26]. Compounds were diluted in DMSO and then in culture medium to achieve a final DMSO concentration of 0.1%. The assay was performed with a 30’ pre-incubation of sporozoites prior to overlaying them on the hepatocytes. In a second mode, compound was added only after sporozoite invasion (3 hr incubation with sporozoite followed by a wash). In both modes, compound containing media were refreshed daily until fixation and analyses at day 4 post invasion.

### *Plasmodium falciparum* transmission assays

Transmission of malaria parasites was studied by Standard Membrane Feeding Assays using luminescent reporter parasite NF54-HGL as described previously [31]. Following incubation of stage V gametocytes with compound for 48 hours a small aliquot was taken for analyses of exflagellation events by light microscopy. The remaining culture material was used to infect *Anopheles stephensi* mosquitoes. Eight days post infection, oocyst intensity was determined by luciferase assays on whole mosquito homogenates as described previously [31]. Gametocyte viability was assessed as described previously [26].

### *Plasmodium falciparum* stage specificity assay

*P. falciparum* NF54-nanoGlo parasites [32] were synchronized by sorbitol treatment as described previously [33] and seeded at a density of 2% parasitemia in 5% hematocrit in RPMI1640 medium supplemented with 10% human type A serum, 367 µM hypoxanthine, 25 mM Hepes and 25 mM sodium bicarbonate and cultured in a semi-automated suspension culture system as described before [34]. Cultures were treated with saturating amount of test compound (5 µM KNX-115), control compound dihydroartemisinin (DHA, 1 µM) or vehicle controls (0.1% DMSO) and samples were taken for thin smears and luminescence measurements at regular intervals. Parasitemia and distribution of parasite stages was determined by microscopic examination of Giemsa-stained thin smears. Luminescence measurements were performed as described previously [32].

### *Plasmodium falciparum in vitro* evolution assays

*P. falciparum* NF54-nanoGlo parasites [32] were seeded at a density of 10^8^ parasites in 10 mL culture medium at 5% hematocrit and cultured in a semi-automated culture system [34] under pressure of KNX-115 at 3xIC_50_. Compound containing medium was refreshed twice per day and parasitemia was followed over time by virtue of Nanoluciferase activity. When luminescence increased, indicative of recrudescent parasites, cultures were passaged at 0.1% parasitemia as normally and drug sensitivity was assessed in asexual blood stage assays.

### Plasmodium falciparum asexual blood stage activity against Dd2 (MRA-156 from BEI-Resources, USA), a clone originally derived from the clinical isolate Indochina III/CDC, and a series of resistant laboratory strains derived from Dd2

The activity was determined with the modified [^3^H]-hypoxanthine incorporation assay, as previously reported [35, 36]

| Strain generated<sup>a</sup> | Mutated locus | Mutations (amino acid changes & references) |
| --- | --- | --- |
| Dd2 | wt | - |
| Dd2 DSM265 | Pfdhodh | [37] |
| Dd2 ELQ-300 | PfcytB | I22L, [38] |
| Dd2 DDD498 | PfeEF2 | Y186N, [39] |
| Dd2 KAF156 | Pfcarl | I1139K, [40] |
| Dd2 MMV048 | Pfpi4k | S743T, [41] |

---

**Figure S1:**
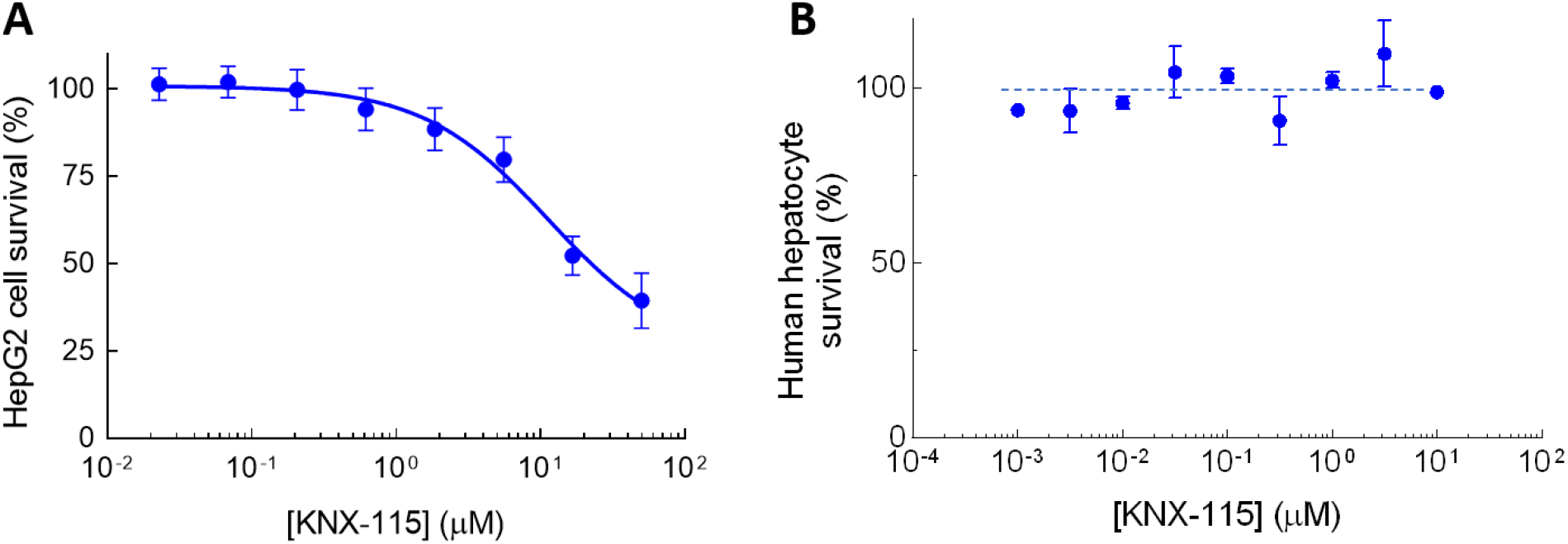
KNX-115 has negligible toxicity in liver cells. A) KNX-115’s toxicity profile on Hep-G2 cells. KNX-115 affects cell viability only at higher concentrations with an EC_50_ >11 µM. Data represents 22 technical replicates from at least 2 biological replicates. B) KNX-115’s liver toxicity profile on primary human hepatocytes. No toxicity was observed even at high concentrations of KNX-115. Data are from two independent experiments each with 2 technical replicates, n = 4.

**Figure S2:**
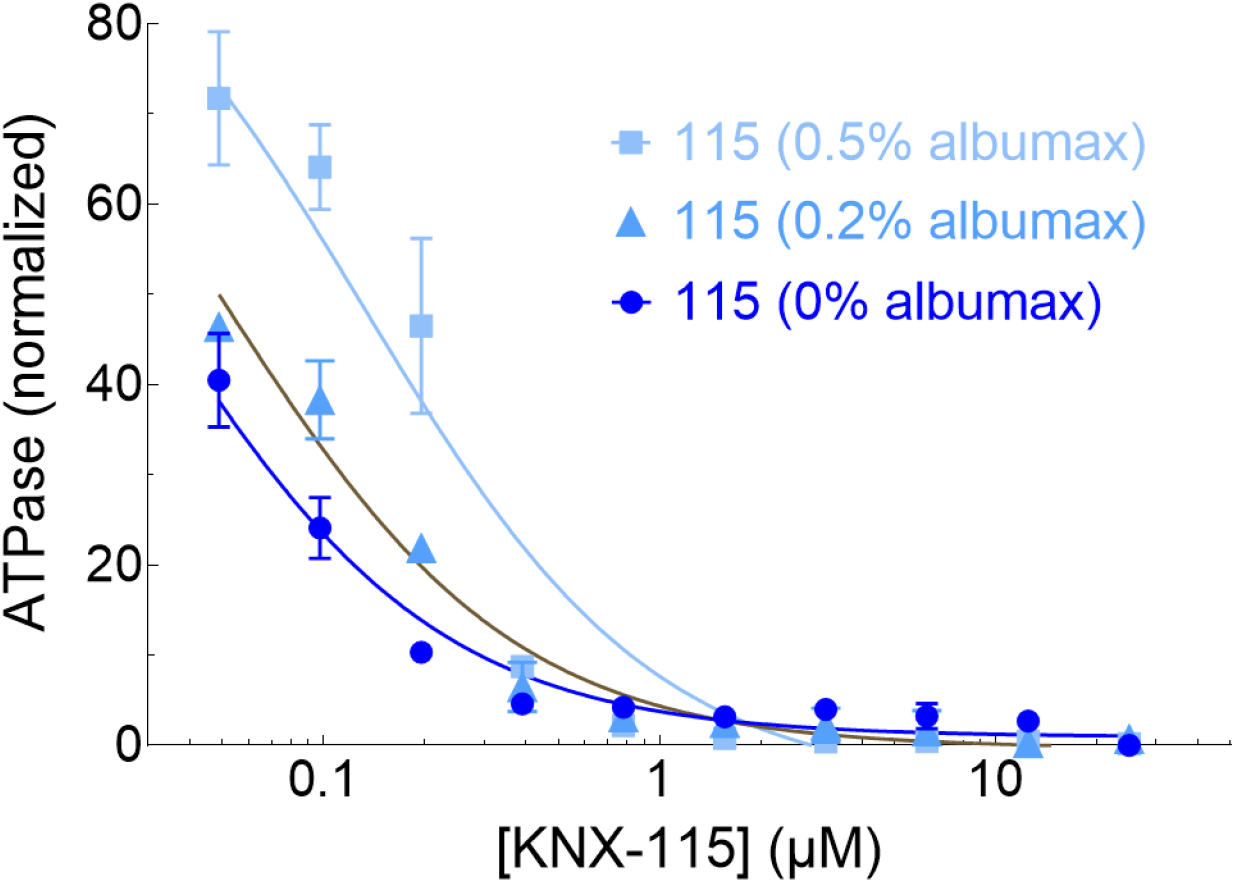
Potency of KNX-115 is weakened in the presence of Abumax. Actin-activated PfMyoA ATPase inhibition by KNX-115 in the presence of 0%, 0.2% and 0.5% Albumax II. Albumax weakens the apparent inhibition potential of KNX-115 in a concentration dependent manner. There is a 3 to 5-fold apparent weakening in the IC_50._ Data are expressed as mean ± SD.

## Notes

### Competing Interest Statement

Darshan V. Trivedi, Annamma Spudich, Kathleen Ruppel, Suman Nag, and James A. Spudich are co-founders of Kainomyx and own stock in the company. Anastasia Karabina, Alice Racca, Heba Wander, Seongwon Jung, and Dan Song are employees of Kainomyx and own stock in the company. Paul V. Ruijgrok and Gustave Bergnes own stock in Kainomyx.

## References

1. Sweeney HL, Holzbaur ELF: Motor Proteins. Cold Spring Harb Perspect Biol 2018, 10.

2. Day SM, Tardiff JC, Ostap EM: Myosin modulators: emerging approaches for the treatment of cardiomyopathies and heart failure. J Clin Invest 2022, 132.

3. Kristeleit R, Evans J, Molife LR, Tunariu N, Shaw H, Slater S, Haris NRM, Brown NF, Forster MD, Diamantis N, et al: Phase 1/2a trial of intravenous BAL101553, a novel controller of the spindle assembly checkpoint, in advanced solid tumours. Br J Cancer 2020, 123:1360–1369.

4. Fabi A, Terrenato I, Vidiri A, Villani V, Tanzilli A, Airoldi M, Pedani F, Magri V, Palleschi M, Donadio M, et al: Eribulin in brain metastases of breast cancer: outcomes of the EBRAIM prospective observational trial. Future Oncol 2021, 17:3445–3456.

5. Chin AC, Day SM: Myosin modulators move forward with FDA approval of mavacamten. Nature Cardiovascular Research 2022, 1:595–596.

6. WHO: World Malaria Report 2021. 2021.

7. Rasmussen C, Alonso P, Ringwald P: Current and emerging strategies to combat antimalarial resistance. Expert Rev Anti Infect Ther 2022, 20:353–372.

8. Cowell AN, Winzeler EA: Advances in omics-based methods to identify novel targets for malaria and other parasitic protozoan infections. Genome Med 2019, 11:63.

9. Bushell E, Gomes AR, Sanderson T, Anar B, Girling G, Herd C, Metcalf T, Modrzynska K, Schwach F, Martin RE, et al: Functional Profiling of a Plasmodium Genome Reveals an Abundance of Essential Genes. Cell 2017, 170:260–272 e268.

10. Wall RJ, Zeeshan M, Katris NJ, Limenitakis R, Rea E, Stock J, Brady D, Waller RF, Holder AA, Tewari R: Systematic analysis of Plasmodium myosins reveals differential expression, localisation, and function in invasive and proliferative parasite stages. Cell Microbiol 2019, 21:e13082.

11. Trivedi DV, Nag S, Spudich A, Ruppel KM, Spudich JA: The Myosin Family of Mechanoenzymes: From Mechanisms to Therapeutic Approaches. Annu Rev Biochem 2020, 89:667–693.

12. Robert-Paganin J, Robblee JP, Auguin D, Blake TCA, Bookwalter CS, Krementsova EB, Moussaoui D, Previs MJ, Jousset G, Baum J, et al: Plasmodium myosin A drives parasite invasion by an atypical force generating mechanism. Nat Commun 2019, 10:3286.

13. Blake TCA, Haase S, Baum J: Actomyosin forces and the energetics of red blood cell invasion by the malaria parasite Plasmodium falciparum. PLoS Pathog 2020, 16:e1009007.

14. Ripp J, Smyrnakou X, Neuhoff MT, Hentzschel F, Frischknecht F: Phosphorylation of myosin A regulates gliding motility and is essential for Plasmodium transmission. EMBO Rep 2022, 23:e54857.

15. Yahata K, Hart MN, Davies H, Asada M, Wassmer SC, Templeton TJ, Treeck M, Moon RW, Kaneko O: Gliding motility of Plasmodium merozoites. Proc Natl Acad Sci U S A 2021, 118.

16. Perrin AJ, Collins CR, Russell MRG, Collinson LM, Baker DA, Blackman MJ: The Actinomyosin Motor Drives Malaria Parasite Red Blood Cell Invasion but Not Egress. mBio 2018, 9.

17. Bookwalter CS, Tay CL, McCrorie R, Previs MJ, Lu H, Krementsova EB, Fagnant PM, Baum J, Trybus KM: Reconstitution of the core of the malaria parasite glideosome with recombinant Plasmodium class XIV myosin A and Plasmodium actin. J Biol Chem 2017, 292:19290–19303.

18. Robert-Paganin J, Xu XP, Swift MF, Auguin D, Robblee JP, Lu H, Fagnant PM, Krementsova EB, Trybus KM, Houdusse A, et al: The actomyosin interface contains an evolutionary conserved core and an ancillary interface involved in specificity. Nat Commun 2021, 12:1892.

19. Moussaoui D, Robblee JP, Auguin D, Krementsova EB, Haase S, Blake TCA, Baum J, Robert-Paganin J, Trybus KM, Houdusse A: Full-length Plasmodium falciparum myosin A and essential light chain PfELC structures provide new anti-malarial targets. Elife 2020, 9.

20. Vahokoski J, Calder LJ, Lopez AJ, Molloy JE, Kursula I, Rosenthal PB: High-resolution structures of malaria parasite actomyosin and actin filaments. PLoS Pathog 2022, 18:e1010408.

21. Williams JW, Morrison JF: [17] The kinetics of reversible tight-binding inhibition. In Methods in Enzymology. Volume 63: Academic Press; 1979: 437–467

22. Morano AA, Dvorin JD: The Ringleaders: Understanding the Apicomplexan Basal Complex Through Comparison to Established Contractile Ring Systems. Front Cell Infect Microbiol 2021, 11:656976.

23. Paton DG, Childs LM, Itoe MA, Holmdahl IE, Buckee CO, Catteruccia F: Exposing Anopheles mosquitoes to antimalarials blocks Plasmodium parasite transmission. Nature 2019, 567:239–243.

24. Paton DG, Probst AS, Ma E, Adams KL, Shaw WR, Singh N, Bopp S, Volkman SK, Hien DFS, Pare PSL, et al: Using an antimalarial in mosquitoes overcomes Anopheles and Plasmodium resistance to malaria control strategies. PLoS Pathog 2022, 18:e1010609.

25. Swann J, Corey V, Scherer CA, Kato N, Comer E, Maetani M, Antonova-Koch Y, Reimer C, Gagaring K, Ibanez M, et al: High-Throughput Luciferase-Based Assay for the Discovery of Therapeutics That Prevent Malaria. ACS Infect Dis 2016, 2:281–293.

26. Schalkwijk J, Allman EL, Jansen PAM, de Vries LE, Verhoef JMJ, Jackowski S, Botman PNM, Beuckens-Schortinghuis CA, Koolen KMJ, Bolscher JM, et al: Antimalarial pantothenamide metabolites target acetyl-coenzyme A biosynthesis in Plasmodium falciparum. Sci Transl Med 2019, 11.

27. Margossian SS, Lowey S: Preparation of myosin and its subfragments from rabbit skeletal muscle. Methods Enzymol 1982, 85 Pt B:55–71.

28. Ikebe M, Hartshorne DJ: Effects of Ca2+ on the conformation and enzymatic activity of smooth muscle myosin. J Biol Chem 1985, 260:13146–13153.

29. Pardee JD, Spudich JA: Purification of muscle actin. Methods Enzymol 1982, 85 Pt B:164–181.

30. Boes A, Spiegel H, Kastilan R, Bethke S, Voepel N, Chudobova I, Bolscher JM, Dechering KJ, Fendel R, Buyel JF, et al: Analysis of the dose-dependent stage-specific in vitro efficacy of a multi-stage malaria vaccine candidate cocktail. Malar J 2016, 15:279.

31. Vos MW, Stone WJ, Koolen KM, van Gemert GJ, van Schaijk B, Leroy D, Sauerwein RW, Bousema T, Dechering KJ: A semi-automated luminescence based standard membrane feeding assay identifies novel small molecules that inhibit transmission of malaria parasites by mosquitoes. Sci Rep 2015, 5:18704.

32. Dechering KJ, Timmerman M, Rensen K, Koolen KMJ, Honarnejad S, Vos MW, Huijs T, Henderson RWM, Chenu E, Laleu B, et al: Replenishing the malaria drug discovery pipeline: Screening and hit evaluation of the MMV Hit Generation Library 1 (HGL1) against asexual blood stage Plasmodium falciparum, using a nano luciferase reporter read-out. SLAS Discov 2022.

33. Lambros C, Vanderberg JP: Synchronization of Plasmodium falciparum erythrocytic stages in culture. The Journal of parasitology 1979, 65 3:418–420.

34. Ponnudurai T, Lensen AH, Meis JF, Meuwissen JH: Synchronization of Plasmodium falciparum gametocytes using an automated suspension culture system. Parasitology 1986, 93 (Pt 2):263–274.

35. Snyder C, Chollet J, Santo-Tomas J, Scheurer C, Wittlin S: In vitro and in vivo interaction of synthetic peroxide RBx11160 (OZ277) with piperaquine in Plasmodium models. Experimental Parasitology 2007, 115:296–300.

36. Chugh M, Scheurer C, Sax S, Bilsland E, van Schalkwyk DA, Wicht KJ, Hofmann N, Sharma A, Bashyam S, Singh S, et al: Identification and deconvolution of cross-resistance signals from antimalarial compounds using multidrug-resistant Plasmodium falciparum strains. Antimicrob Agents Chemother 2015, 59:1110–1118.

37. Phillips MA, Lotharius J, Marsh K, White J, Dayan A, White KL, Njoroge JW, El Mazouni F, Lao Y, Kokkonda S, et al: A long-duration dihydroorotate dehydrogenase inhibitor (DSM265) for prevention and treatment of malaria. Sci Transl Med 2015, 7:296ra111.

38. Stickles AM, de Almeida MJ, Morrisey JM, Sheridan KA, Forquer IP, Nilsen A, Winter RW, Burrows JN, Fidock DA, Vaidya AB, Riscoe MK: Subtle changes in endochin-like quinolone structure alter the site of inhibition within the cytochrome bc1 complex of Plasmodium falciparum. Antimicrob Agents Chemother 2015, 59:1977–1982.

39. Baragana B, Hallyburton I, Lee MC, Norcross NR, Grimaldi R, Otto TD, Proto WR, Blagborough AM, Meister S, Wirjanata G, et al: A novel multiple-stage antimalarial agent that inhibits protein synthesis. Nature 2015, 522:315–320.

40. Meister S, Plouffe DM, Kuhen KL, Bonamy GM, Wu T, Barnes SW, Bopp SE, Borboa R, Bright AT, Che J, et al: Imaging of Plasmodium liver stages to drive next-generation antimalarial drug discovery. Science 2011, 334:1372–1377.

41. Paquet T, Le Manach C, Cabrera DG, Younis Y, Henrich PP, Abraham TS, Lee MCS, Basak R, Ghidelli-Disse S, Lafuente-Monasterio MJ, et al: Antimalarial efficacy of MMV390048, an inhibitor of Plasmodium phosphatidylinositol 4-kinase. Sci Transl Med 2017, 9.

